# PocketBagger: Generalizable pocket druggability prediction via positive–unlabeled learning

**DOI:** 10.64898/2026.05.15.725505

**Authors:** Phillip W Gingrich, Ansuman Biswas, Ioan L Mica, Kevin M Brammer, Zhigang Shu, David S Maxwell, Kaitlyn P Russell, Bissan Al-lazikani

## Abstract

Reliable structure-based prediction of small-molecule druggability is hindered by a fundamental labeling problem. Experimentally confirmed liganded sites (positives) are observable, but credible “undruggable” pockets (negatives) are almost impossible to define. Standard supervised machine learning consequently relies on arbitrary definitions of ‘undruggable’, leading to bias and false negatives. Here we introduce PocketBagger, a positive–unlabeled (PU) learning framework for pocket druggability prediction trained exclusively on experimentally determined Protein Data Bank^1^ (PDB) structures. PocketBagger uses PU bagging to learn key features associated with reliable ‘druggable’ pockets and considers all remaining pockets in the structurally characterized proteome as unlabeled. We demonstrate the capability of PocketBagger through the training of a simple Random Forest classifier and demonstrate its power in recall (0.804), even when challenged with increasingly difficult generalizability assessments and entire protein-family hold outs. We benchmark and demonstrate the added value of PU learning by comparing PocketBagger to a leading deep-learning predictor. However, PocketBagger is intended to be used as a framework for any model architecture. Along with the code, the data generated by PocketBagger are deployed in canSAR.ai, providing scalable, generalizable pocket druggability predictions to the drug discovery community.

## Introduction

Drug discovery can be a decades-long and expensive process, often costing more than $2.8 billion per approved drug and resulting in high failure rate of candidate molecules during clinical trials(∼90%)^2^. Among the earliest and most crucial steps in drug discovery is target selection.

Small molecule ‘druggability’ was described >20 years ago and used to define validated drug targets and their homologues. First estimates of a ‘druggable genome’ in 2002 were conservative and relied on homology to previously drugged, ‘privileged’ families^3^. Since then, experimental and computational technology^4^ have been employed for extending beyond these. Some methods attempt to enhance this similarity beyond protein sequence homology using e.g. vectorized pocket properties (“PocketVec”)^5^. Most approaches, however, have opted to employ machine learning (ML) algorithms to learn properties of druggable pockets by (1) defining ‘pockets’ using tools such as SURFNET^6^ or Fpocket^7-9^), and, then, (2) predicting their druggability using a classification algorithm (e.g. canSAR^10, 11^, DrugMiner^12^). Whether it’s traditional ML or increasingly sophisticated deep learning approaches such as GrASP^13^, all methods rely on the same assumptions for the initial training set: that pockets can be reliably defined as either druggable (positive) or ‘undruggable’ (negative). While ‘druggable’ pockets can be defined as those experimentally shown to bind a drug or drug-like molecule^10^, empirically defining ‘undruggable’ pockets is impossible. This is a result of how little of the human proteome has been chemically explored^14, 15^—let alone with “drug-like” molecules. This is highlighted by the fact that across PDB structures in canSAR, only 6.6% of pockets contain any ligand. It is therefore a naïve assumption that unligandable pockets are always negative ‘undruggable’ pockets. Moreover, since experimentally validated druggable sites are so few, generating independent test sets to appropriately assess generalizability beyond what is currently known is extremely challenging. As a result of the above limitations, current methods remain biased and constrained to varying degrees of established druggability and skewed against truly novel druggable sites.

To address this limitation, we propose employing the positive–unlabeled (PU) learning framework for pocket druggability prediction. In PU learning, the training set contains a small set of confirmed positives and a large unlabeled pool that is a mixture of both positives and negatives^16^. Critically, the unlabeled class is treated as *unknown* rather than assumed to be negative. PU bagging^17^ is a suitable PU approach as it avoids committing any unlabeled example to a definitive negative label by repeatedly training classifiers on all positives and different resampled subsets of the unlabeled pool, aggregating predictions across bags. Given that “undruggable” pockets are rarely observed with the same confidence as druggable ones, as illustrated by historically “undruggable” targets later shown to be tractable (e.g., KRAS), PU learning provides a more appropriate learning paradigm for pocket druggability than standard supervised classification.

In this work, we introduce PocketBagger, a positive–unlabeled learning approach for structure-based pocket druggability prediction trained exclusively on experimentally determined PDB structures. PocketBagger uses PU bagging to learn from pockets labeled as positive by the presence of curated drug-like ligands while treating the remaining pockets as unlabeled, avoiding reliance on unreliable “undruggable” labels. Generalizability is a primary design aim throughout; hyperparameter selection and performance evaluation are conducted with sequence-cluster– aware validation to prevent homology leakage, complemented by chain-aware and protein-aware splits, targeted protein-family holdouts, and label permutation controls to quantify robustness under increasingly strict out-of-distribution settings. Finally, we benchmark PocketBagger against a contemporary deep learning-based site predictor (GrASP) using a strict temporal and novelty-based holdout set of post-publication structures drawn from previously unseen sequence clusters, providing a forward-looking comparison on independently verifiable experimental structures. This comparison is not made for the sake of competition; rather, we juxtapose the benefit of a PU approach even when applied to a simpler ML method with the leading deep learning predictor as the ‘yard stick’ for performance.

## Results and Discussion

### Improved training of druggability labeling with PU learning

Herein, we use the term “druggability” for simplicity and define druggability of a pocket as that pocket’s compatibility with binding a “drug-like” small molecule with properties generally consistent with Lipinski^18^, Veber^19^, and Ghose^20^ filters, as indicated in our Methods. For this study, we make no assumptions about functional or therapeutic impact.

We predicted druggability using the PocketBagger workflow as shown in Fig. 1. Briefly, we trained a positive– unlabeled bagging classifier on SURFNET^6^-derived pocket descriptors from experimentally determined PDB structures, defining positive pockets by the presence of curated drug-like and “reasonable” ligands (Methods). After sequence-cluster assignment using MMSeqs2^21^, 580,304 unique protein chains spanning 24,101 sequence-based clusters were carried forward for model development. Pocket geometry and physicochemical context are highly correlated within homologous proteins. This significantly risks overinflating performance with limited generalizability to unseen protein classes. To address this, we prioritized generalizability in both model selection and evaluation by using sequence-based clusters as groups to drive group-aware splitting and investigating performance across increasingly stringent split strategies (defined in Methods as random, chain-aware, protein-aware, and cluster-aware), supplemented further by targeted family holdouts and label permutation controls. This evaluation design enables a realistic estimate of performance on novel proteins and provides a foundation for benchmarking PocketBagger against contemporary site predictors.

**Figure 1:**
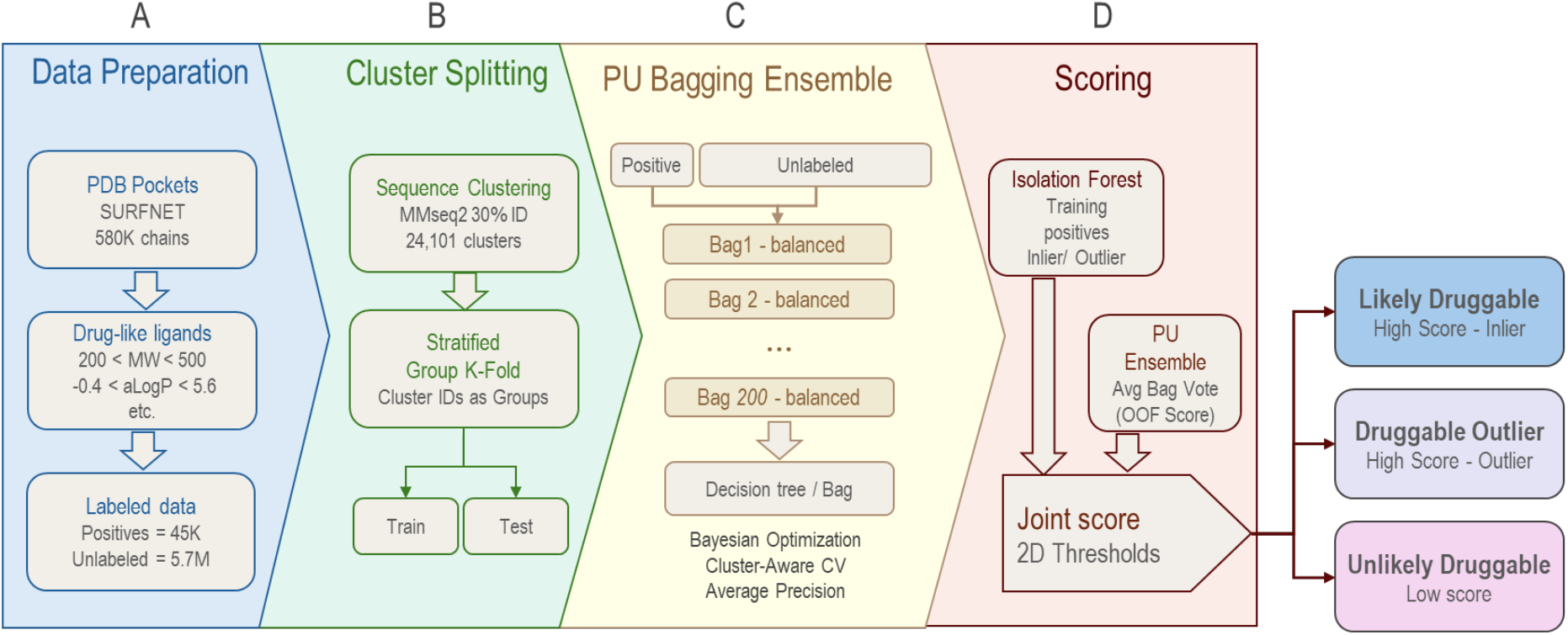
Overview of the PocketBagger workflow. A) Data curation involved querying pockets within canSAR.ai and labeling pockets according to the presence of “drug-like” ligands in the pocket. The complete list of physicochemical properties used is outlined in the Methods section. B) Structures are clustered using MMSeqs2 to drive cluster-aware splitting for holdout sets. C) A PU Bagging ensemble of decision tree classifiers is tuned using cluster-aware splitting and finally trained. D) An isolation forest trained on training positives and the ensemble of classifiers jointly score records, enabling further segmentation of predictions. On the training data, out of fold (OOF) scores are generated to facilitate unbiased data mining.

## Model Performance

### Generalization under increasingly stringent evaluation splits

We first quantified PocketBagger’s performance under a hierarchy of nested cross-validation (CV) splits designed to progressively reduce information leakage from structurally or evolutionarily related proteins (Fig. 2A & 2B). Under a standard random split, PocketBagger achieved high true-positive recall (0.895 ± 0.003, mean ± s.d., *n* = 5 repeats), and highly ranked held out true positives as indicated by recall@K_i_ (0.948 ± 0.002, mean ± s.d., *n* = 5 repeats). However, as described above, generalizability needs to be tested under realistic conditions. As expected for structure-derived pocket descriptors, performance depends on whether related structures can appear in both training and test folds. Enforcing chain-aware splitting (i.e., all pockets from a given PDB chain confined to a single fold) preserved overall recall (0.895 ± 0.005) but reduced structure-level recovery as captured by recall@K_i_ (0.870 ± 0.005, Fig. 2A). When the split was made more stringent by holding out entire UniProt identifiers (protein-aware), recall decreased to 0.812 ± 0.042 and recall@K_i_ to 0.815 ± 0.033, consistent with a larger shift in the distribution of test proteins relative to training. While it is difficult to make like-to-like comparisons (as detailed in the supplemental information), the most comparative metric from GrASP is DCA Recall at Top N between 0.775 and 0.813, showing a highly comparable level of performance under similar splitting strategies^13^. A more strict and realistic regime—sequence cluster–aware splitting at 30% identity to remove related homologues—had little impact on recall (0.804 ± 0.055 and recall@K_i_ to 0.805 ± 0.059). This approach limits data leakage caused by homology, providing an unbiased estimate of generalizability, extending beyond simple protein similarity.

**Figure 2.**
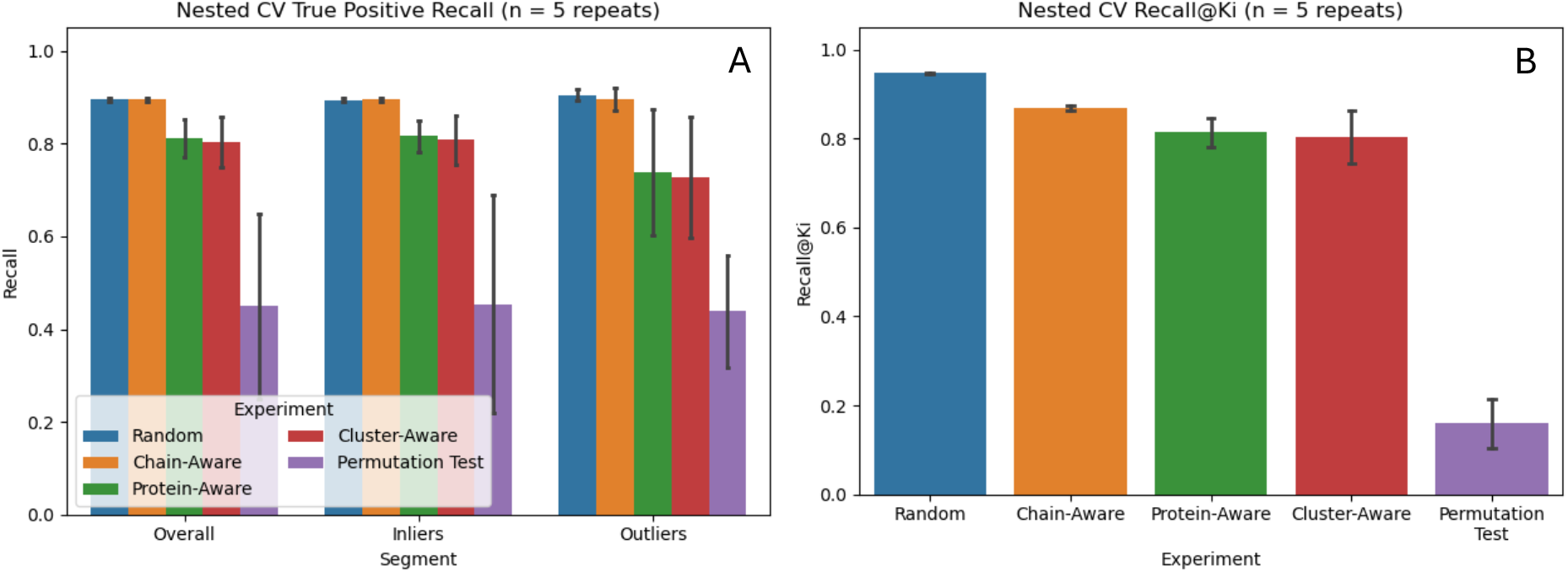
Nested cross validation performance under varying splitting strategies. A) Recall values indicate robust but decreasing performance consistent with increasingly stringent splitting strategies to prevent leakage between train and test splits. B) Recall@K_i_ values similarly indicate PocketBagger’s ability to highly rank known positives in holdout sets with expected performance drops and increased variability under rigorous splits. By both measures, the permutation test provides a null baseline for comparison. Values reported represent means, and error bars indicate the standard deviation.

In a real-world setting, new data from novel protein structures may have pockets with feature distributions that are different to the training data. To probe such a distribution shift, we additionally stratified test pockets into “inliers” and “outliers” relative to the training positives using an isolation forest^22, 23^ trained on training-set positives for each split (Fig. 2A). Across split types, inlier recall closely tracked overall recall (e.g., 0.808 ± 0.054 under cluster-aware splitting), whereas outlier performance was generally lower and more variable (cluster-aware outliers: 0.727 ± 0.130). This pattern supports the intended role of the anomaly detection model: pockets that fall farther from the positive training manifold represent a more out-of-distribution setting. Such records are more likely to correspond to genuinely novel pocket chemistries or atypical structural contexts, and labeling them with the auxiliary isolation forest model affords additional nuance for researchers to consider in investigating such a site.

Finally, a label permutation test confirmed that the measured performance is not attributable to trivial dataset artifacts. Under permuted labels, overall recall dropped to 0.450 ± 0.200 and recall@K_i_ to 0.159 ± 0.055 (Fig. 2A–B), providing a conservative null baseline for comparison. This comparison highlights significant improvements to recall and recall@K_i_ over the random setting.

### Targeted protein-family holdouts stress-test for cross-family generalization

The PDB is enriched with historically “privileged” and highly studied drug-target families. The sequence clustering above significantly reduced family-based contamination and eliminated leakage through obvious homology. However, the most challenging test would be to examine how PocketBagger performs on entirely novel and previously unseen protein folds. To test this, we devised a targeted family holdout test to mimic this challenging real-world use case. We focused on three families that are both drug-discovery–relevant and strongly represented in the structural records— namely protein kinases, nuclear hormone receptors, and G protein–coupled receptors—and treated each family as an explicit test partition. To ensure this was a true generalization test rather than a near-duplicate test, pockets in structures belonging to sequence-based clusters found in the heldout family were additionally removed from the training set, further preventing data leakage when the split was defined at the UniProt-family level. This removal of overlapping clusters was explicitly applied for the family holdouts. Pocket-level true-positive recall remained high overall for protein kinases (0.907) and G protein–coupled receptors (0.894) and was highest for nuclear hormone receptors (0.960) (Fig. 3A). Performance remained strong for structure-level ranking and retrieval (recall@K_i_), achieving 0.909 for protein kinases, 0.938 for nuclear hormone receptors, and 0.838 for G protein– coupled receptors (Fig. 3B). Performance under stratification by inlier/outlier status (relative to the training-positive manifold) varied compared to the cluster-aware performance reported above. Inliers remained consistently strong across families (≈0.91–0.96), whereas outlier recall was family-dependent, dropping dramatically for protein kinases (0.308) and increasing slightly for nuclear hormone receptors (0.981). Rigorously explaining this variability is difficult given the small number of outliers in each family holdout test (50 for protein kinases, 52 for nuclear hormone receptors, and 83 for GPCRs). Nevertheless, segmentation by the isolation forest is useful as it flags outliers. Such pockets whose feature profiles are less supported by the positive training distribution are called to attention so that researchers can readily identify records likely in need of expert inspection.

**Figure 3.**
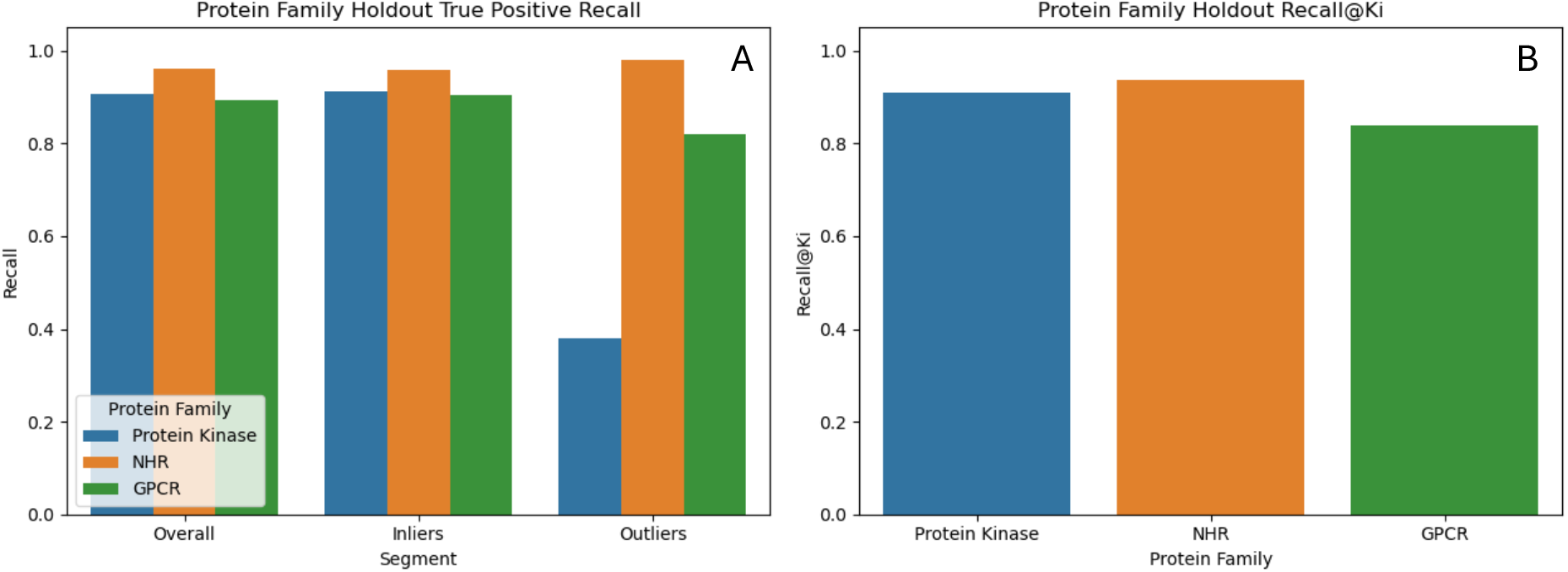
Targeted protein family holdout performance. A) Recall values indicate PocketBagger’s ability to recall known positives despite these families and similar structures being withheld from training. B) Recall@K_i_ values highlight PocketBagger’s ability to highly rank known druggable pockets in protein family structures. By both measures, generalizability of the approach is demonstrated by novelty splits retrospectively.

Collectively, these targeted holdouts complement the random/chain/protein/cluster-aware tests by demonstrating that PocketBagger’s generalizability to recall and highly rank druggable pockets is preserved when entire, drug-relevant protein families are withheld from training.

### Temporal and novelty-based benchmark against GrASP

As we previously noted, direct comparisons between methods are challenging. To make a more direct comparison on a strict external benchmark, we compared PocketBagger against GrASP using a small but challenging holdout set comprising of 26 PDB chains released after July 26, 2023, belonging to clusters not observed before that date. This date was chosen as GrASP model weights were deposited to GitHub on that same date.

Because GrASP outputs atom-level probabilities, we converted predictions to a site-level score anchored to the observed ligand by aggregating probabilities over protein heavy atoms within 4 Å of the bound drug-like ligand, using both the mean and the maximum probability in that neighborhood, and applying GrASP’s recommended site-call threshold of 0.3.

Notwithstanding the small sample size, PocketBagger’s recall on this holdout (Fig. 4) was consistent with its performance under the cluster-aware evaluation regime, supporting that the nested, leakage-controlled performance estimates translate to realistic production settings. Moreover, PocketBagger exceeded GrASP under both mean- and max-aggregated site calling on this forward-looking benchmark, indicating that a PU-bagging approach trained on experimentally observed positives can match and, in this setting, outperform a contemporary deep learning–based site predictor under simultaneous temporal and novelty constraints. We emphasize that *n* = 26 chains is not intended to support fine-grained statistical claims; rather, this set functions as a prospective-style stress test to exemplify the generalizability of our PU approach. We assert this generalizability highlights the utility of the PU framework and suggest wider adoption where applicable alongside classical or deep learning methodologies.

**Figure 4.**
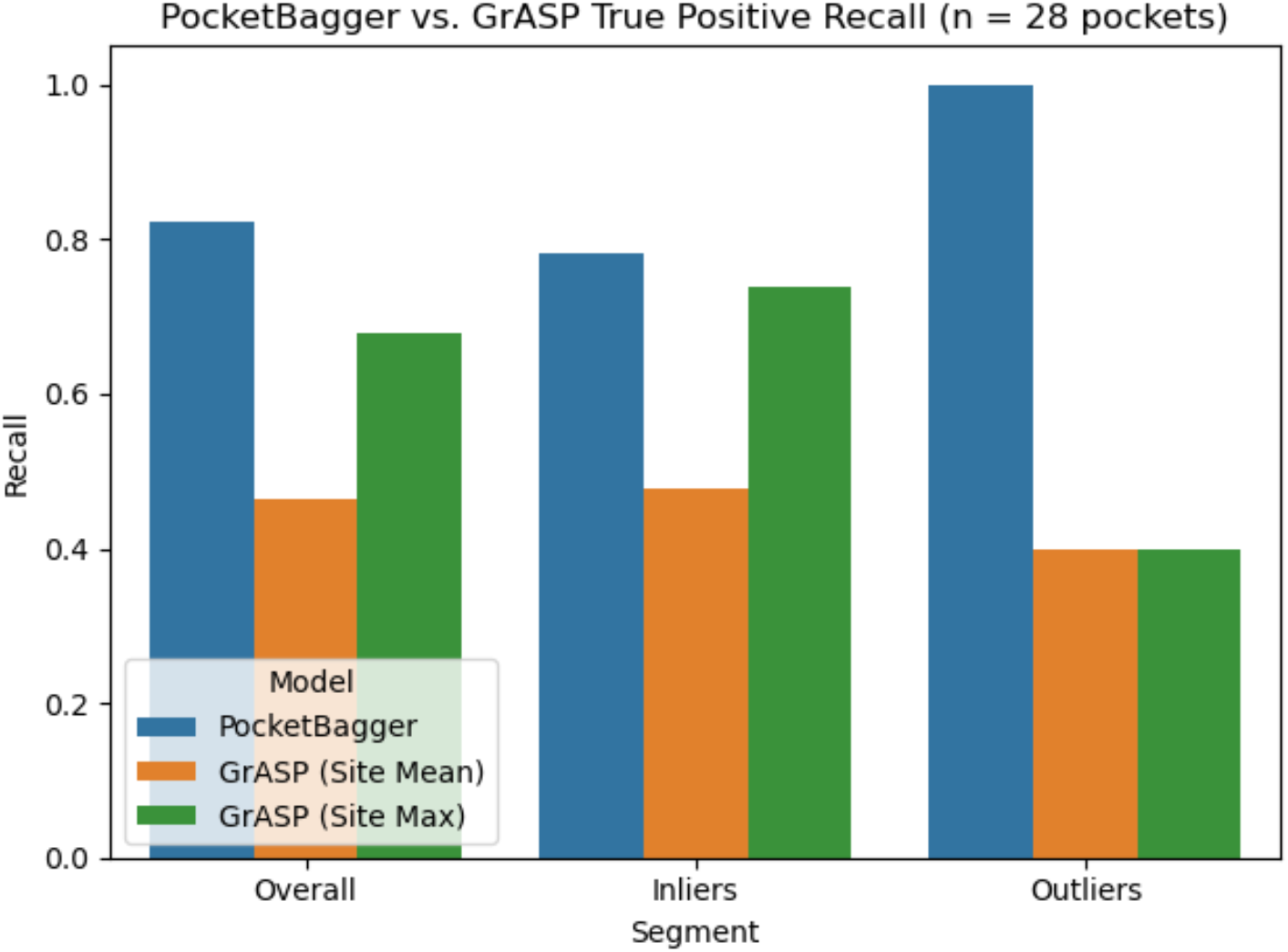
Temporal and novelty based split comparative performance between PocketBagger and GrASP. A dataset of 26 structures published after July 26, 2023 that do not belong to a sequence-based cluster observed before that date was mined from the PDB, with pocket data pulled from canSAR.ai. The 26 structures contained 28 pockets occupied by a “drug-like” ligand. By both site mean and maximum probabilities used from GrASP, PocketBagger’s recall exceeds GrASP, to include across inlier and outlier segments.

**Figure 5.**
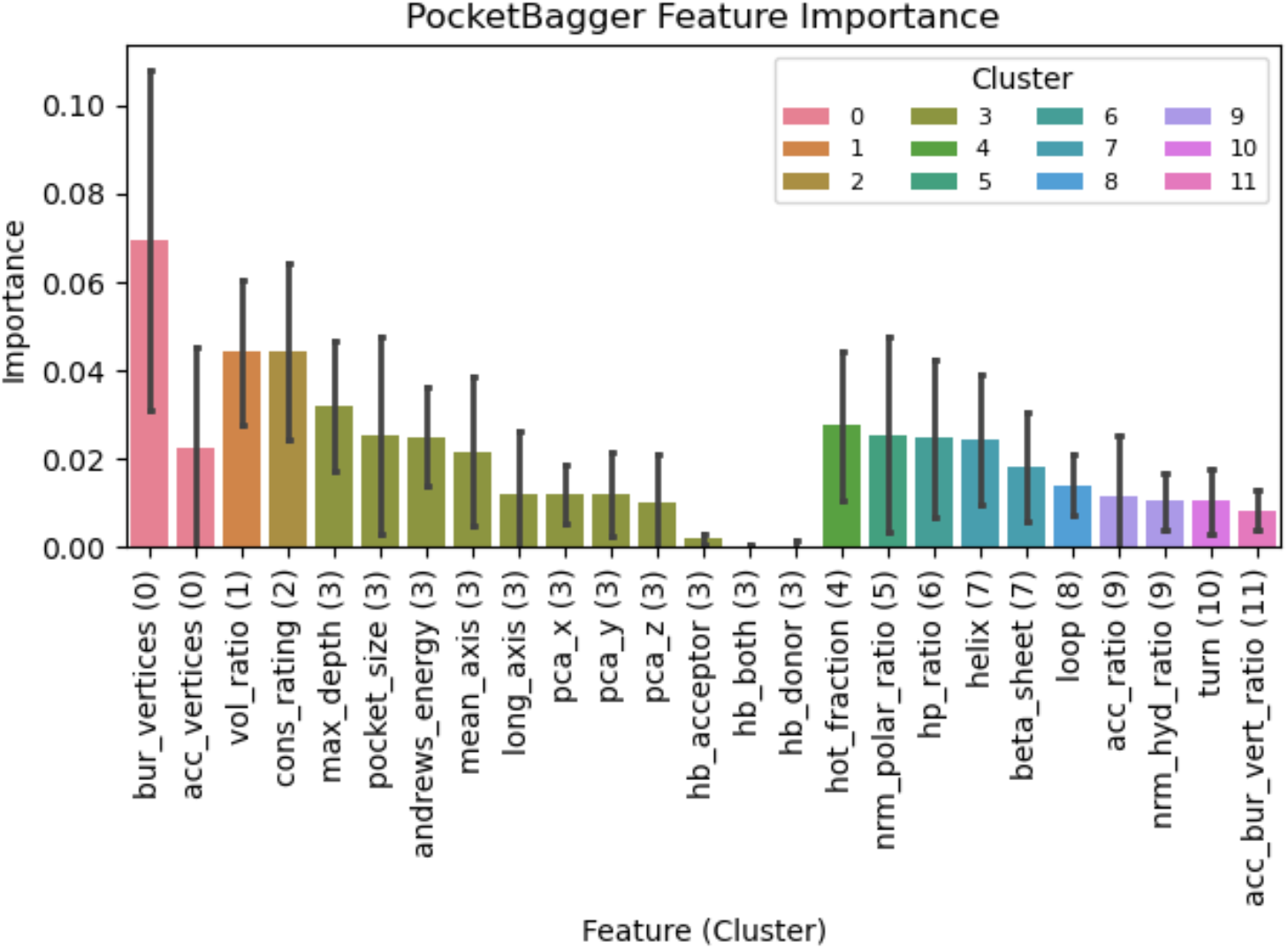
Permutation feature importance for PocketBagger. Importances were computed using average precision as the driving metric. High observed importances in buriedness, pocket volume ratio, roughness (as measured by cons_rating), depth, and pocket volume conceptually match properties generally understood to influence druggability. Values reported represent means across stratified group-aware 5-fold splits, and error bars indicate the standard deviation.

### Feature Importance Analysis

Permutation-importance analysis, aggregated five stratified group-aware folds, indicates that PocketBagger’s predictions are driven primarily by geometric enclosure, pocket size, roughness, and shape descriptors, rather than any single idiosyncratic feature. The most influential variables include buried vertices (bur_vertices) and related measures of enclosure, volume ratio (vol_ratio) and overall size/extent descriptors (e.g., pocket_size, max_depth, mean_axis), together with cons_rating, a measure of pocket surface roughness (i.e., complexity of surface peaks and valleys). Consistent with ligandability intuition, these signals collectively capture whether a cavity is sufficiently enclosed, voluminous, deep, and structurally articulated to support persistent protein–ligand contacts. The correlation-clustering of features shows that importance is distributed across coherent feature groups rather than being dominated by redundant correlates. Secondary contributions from physicochemical composition (e.g., nrm_polar_ratio, hp_ratio) and local “hot spot” proxies (hot_fraction) suggest that PocketBagger further refines predictions based on interaction potential once basic geometric tractability is satisfied. Overall, the importance profile supports that PocketBagger learns interpretable decision rules centered on pocket geometry and roughness first, with composition acting as a second-order discriminator across the broader set of unlabeled cavities.

#### Score- and anomaly-informed thresholding of unlabeled pockets

To inform prospective thresholding (i.e., deciding where to “draw the line” when promoting unlabeled pockets to likely positives for future datasets) we combined three complementary cutoffs in the joint space of out-of-fold (OOF) PocketBagger scores and isolation-forest anomaly scores (Fig. 6). First, we applied upper Tukey-fence thresholds to the distribution of scores in unlabeled samples at two stringencies (k = 1.5 and k = 3.0), providing pragmatic “high-confidence” score cutoffs for prioritization. Second, we fit an Elkan–Noto^24^ (PU) class-prior model with isotonic calibration and used it to derive a probability threshold corresponding to the estimated positive class-prior. To reduce bias from overrepresented protein families and partially mitigate violations of the SCAR assumption in training the Elkan–Noto classifier, we weighted training examples by the inverse size of their sequence cluster and when generating group-aware out-of-fold probabilities. Third, we trained an isolation forest on the training positives to quantify how “positive-like” each pocket is in feature space and used a balanced random forest to empirically determine an isolation-score boundary separating the positive manifold from the broader unlabeled background. Pockets falling into the dense positive region of the 2D score–anomaly histogram are reasonably separable from the dense unlabeled region using these boundaries.

**Figure 6.**
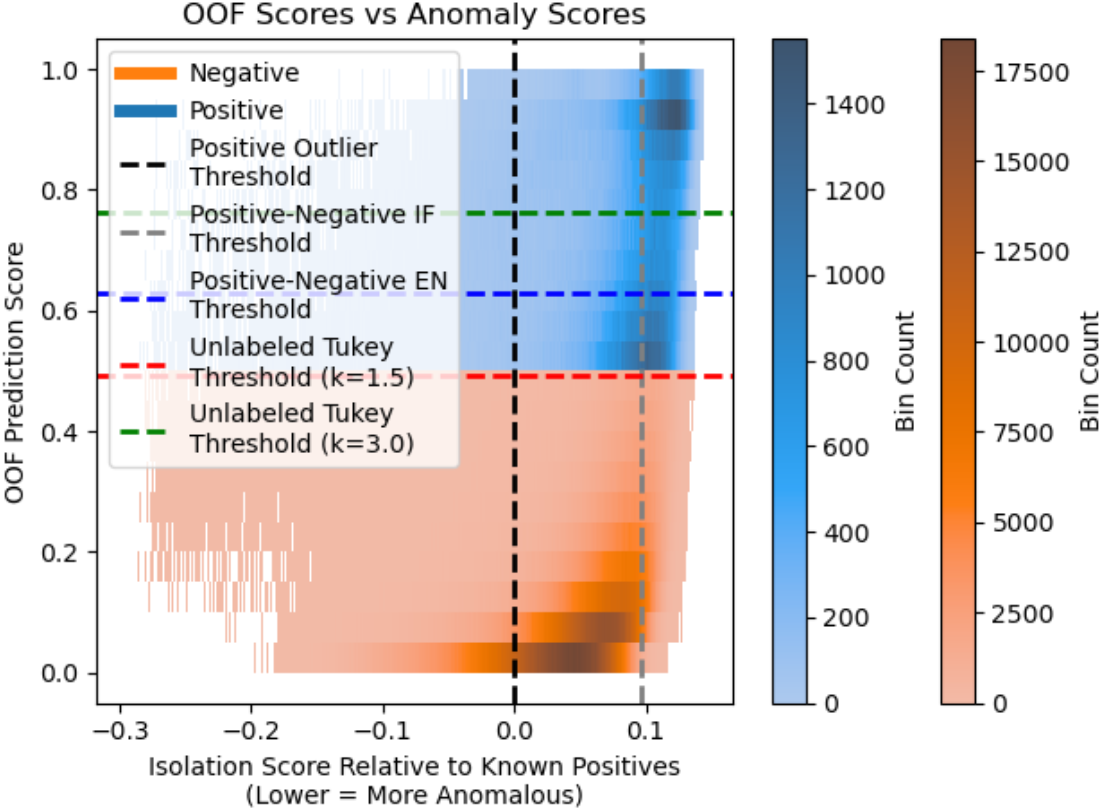
2D histogram of unbiased out of fold (OOF) prediction scores versus isolation forest anomaly scores. Positive predictions were taken as those records with OOF scores of 0.5 or greater. Various thresholds are depicted that allow for systematic segmentation of samples in a production setting to increase confidence in positive classification over a naïve threshold of 0.5 alone.

Importantly, these thresholds serve different roles and are intended to be used in combination. Tukey fences are heuristics that favor precision at the expense of coverage, believing “druggable” pockets are rare given the difficulty with which the community identifies novel tractable sites. The Elkan– Noto threshold provides a prevalence-informed threshold under an explicit (and imperfect) PU assumption to suggest what fraction of the data is potentially positive. The anomaly score helps compliment prediction scores with in- or out-of-distribution assessments so that expert judgement and intuition can be further applied. While the SCAR assumption remains violated in structure-derived druggability labels and no single cutoff is “correct,” together these approaches provide a systematic and auditable way to threshold predictions, increasing confidence (and likely precision) when calling positives from the unlabeled pool in production settings.

### Conclusion

PocketBagger addresses a central challenge in structure-based druggability prediction: the lack of reliable negative (“undruggable”) labels and the resulting bias introduced by standard supervised training. By framing pocket druggability as a positive–unlabeled learning problem and using PU bagging over SURFNET-derived pocket features from experimentally determined PDB structures, PocketBagger learns to prioritize likely ligandable pockets while treating the remaining cavities as unlabeled rather than forcing uncertain negatives.

In a generalizability-first evaluation design—spanning random, chain-aware, protein-aware, and sequence-cluster–aware splits—PocketBagger maintains strong recall as leakage controls increase, providing a realistic estimate of performance on novel proteins and unseen sequence families. These results are reinforced by targeted drug-relevant family holdouts and label-permutation controls. Using a small but deliberately challenging temporal and novelty-based benchmark against a contemporary deep learning site predictor (GrASP), PocketBagger’s performance is consistent with its cluster-aware estimates and exceeds GrASP under prospective conditions despite PocketBagger using a simpler ML framework. These results illustrate that using such a PU framework enhances performance, robustness, and, most importantly, generalizability, adding value to any predictive tool regardless of the ML architecture it ultimately employs. PocketBagger is deployed in canSAR.ai, enabling users to explore protein structures for likely druggable sites using a model explicitly designed and evaluated for generalizability.

## Online Methods

All scripts were generated using Python 3.12, leveraging NumPy, pandas, and scikit-learn for machine learning. While intended for imbalanced setting, imbalanced-learn was used to drive PU bagging. Readers can refer to the GitHub repository for all source code at https://github.com/a3d3a/PocketBagger

### Dataset preparation for small molecules

First, we compiled a curated set of “drug-like” Protein Data Bank (PDB) ligands. All ligand records and their associated isomeric SMILES were downloaded from the wwPDB Components file (https://files.wwpdb.org/pub/pdb/data/monomers/Components-smiles-stereo-oe.smi). Using RDKit^25^, ligands were standardized and physicochemical properties were computed, including molecular weight (MW), calculated LogP (aLogP), topological polar surface area (TPSA), hydrogen-bond donors (HBD) and acceptors (HBA), number of rotatable bonds (ROTBONDS), heavy-atom count (HVY_ATMS), and quantitative estimate of drug-likeness (QED).

A “drug-like” subset was defined using a combined Lipinski-^18^, Veber-^19^, and Ghose-style envelope together with a minimum drug-likeness constraint: QED ≥ 0.5, 200 ≤ MW ≤ 500 Da, -0.4 ≤ aLogP ≤ 5.6, HBA ≤ 10, HBD ≤ 5, TPSA ≤ 140 Å^2^, ROTBONDS ≤ 10, and HVY_ATMS ≥ 10. To exclude crystallographic artifacts and non–small-molecule entities, ligands were additionally required to be organic (contain carbon), composed solely of standard non-metal atoms (H, B, C, N, O, F, Si, P, S, Cl, Se, Br, and I), and not corresponding to canonical or non-canonical amino acids or polymer-like components.

Because hard physicochemical thresholds can exclude ligands that lie near, but not within, the drug-like envelope, we expanded the ligand panel using a data-driven approach. An isolation forest was trained in the same physicochemical descriptor space on the stringent “drug-like” set, and predicted inliers among the initially excluded ligands were recovered as “reasonable” small molecules (again restricted to organic, standard-atom ligands and excluding amino-acid–like entities).

The final ligand panel comprised both the stringent “drug-like” ligands and these near-boundary “reasonable” ligands. Pockets occupied by any ligand from this combined set were labeled as positives (“druggable”) for subsequent positive–unlabeled learning. A PCA was performed (Fig. S1) and example molecules were sampled from each class (Fig. S2), both for the sake of visualization.

Additionally, feature sensitivity in the isolation forest was performed to better understand what features contributed to “drug-likeness” by our approach (Fig. S3). Property distributions of each subset of molecules are also shown (Fig. S4-S20).

### Protein Clustering

All PDB protein chains were retained to preserve maximal structural diversity across the proteome. To control for sequence redundancy and prevent information leakage, we clustered the PDB SEQRES sequences (downloaded from https://files.wwpdb.org/pub/pdb/derived_data/pdb_seqres.txt.gz) at 30% sequence identity using MMseqs2. Each chain was assigned to a sequence-based cluster.

After clustering and mapping within the canSAR pocket dataset, 580,304 unique protein chains were carried forward for model development, grouped into 24,101 sequence-based clusters. These cluster assignments were used in cluster-based group-aware splitting in various experiments to ensure that homologous proteins were not distributed across evaluation sets. Importantly, clusters were not reduced to single representative structures; multiple chains per cluster were retained so that the model could learn conformational and pocket-level variability within protein families, reflecting the heterogeneity encountered in practical structure-based drug discovery.

### PU Bagging Learning Scheme (PocketBagger)

#### Pocket feature dataset and labeling

Pocket-level descriptors were retrieved from the SURFNET-based pocket dataset within canSAR as previously described^26^. Each record corresponds to a single pocket instance and includes geometric, compositional, and structural features (e.g., pocket size; buried and accessible vertex counts; hydrophobic and polar ratios; hydrogen-bond donor/acceptor counts; secondary-structure composition; depth and axis-length measures; and principal component coordinates). These are fully defined in the supplementary text under “Pocket Descriptors”. Records lacking complete feature vectors or valid chain/cluster assignments were removed. Pockets were labeled as positive (“druggable”) if they were occupied by at least one ligand from the curated “drug-like”/reasonable HET panel described above. All remaining pockets were treated as unlabeled.

#### Positive–unlabeled bagging framework

PocketBagger was implemented as a positive–unlabeled (PU) bagging ensemble using the BalancedBaggingClassifier from *imbalanced-learn*, with a decision tree as the base estimator. In each bag, all confirmed positives were retained, while an equally sized subset of unlabeled pockets was treated as provisional negatives. Balanced sampling within each bag avoids consistently assuming all unlabeled pockets are negative.

#### Bayesian hyperparameter optimization with cluster-aware validation

Hyperparameters of the decision-tree base estimator were optimized using Bayesian optimization (BayesSearchCV). We tuned: (i) maximum tree depth (max_depth) and (ii) the number of features considered at each split (max_features). The upper bound for max_depth was set empirically by first fitting an intentionally over-parameterized ensemble and taking the maximum observed tree depth as the ceiling for the Bayesian search.

Hyperparameter selection used average precision (area under the precision–recall curve) as the optimization objective as it is still the expectation that true positives are generally ranked more highly than most unlabeled data points. During tuning, validation folds were enforced with sequence-cluster– aware splitting (cluster-disjoint folds), promoting generalization beyond homologous proteins. Early stopping was applied to terminate Bayesian optimization when no improvement was observed over successive iterations.

#### Isolation forest segmentation of inliers and outliers

For each evaluation split (i.e., for each train/test instance), an isolation forest was trained only on the training-set positives to characterize the region of feature space occupied by confirmed druggable pockets. To reduce overrepresentation of large sequence clusters, positive samples were weighted inversely by the number of positives in their cluster during isolation-forest fitting.

The isolation forest’s contamination parameter was estimated from the distribution of positive isolation scores using a bootstrap Tukey-fence procedure (k = 1.5). Specifically, isolation scores for training positives were bootstrapped repeatedly, and for each bootstrap replicate, a lower Tukey fence was computed. The expected contamination was then taken as the average fraction of bootstrapped scores falling below their replicate-specific fence. This Tukey-derived contamination was used to set the isolation forest’s contamination for that split (with a conservative cap applied when appropriate). Test pockets were then assigned as *inliers* or *outliers* based on the isolation forest decision function threshold, enabling stratified performance reporting by distributional proximity to the training positives.

In the final production isolation forest, feature sensitivity was computed by feature permutation and calculating the difference in the spearman rho between the anomaly scores before and after feature permutation. This was done for 5 repeats using the positive pockets across the dataset. Results are presented in Figure S2.

#### Evaluation regimes

True positive recall was used as the performance metric on all hold-out test sets given only positives labels are known. Model performance was assessed under multiple partitioning schemes of increasing strictness:

1. Random split: stratified k-fold splitting without grouping.
2. Chain-aware split: stratified group k-fold splitting grouped by PDB chain (all pockets from a chain confined to one fold).
3. Protein-aware split: stratified group k-fold splitting grouped by UniProt identifier.
4. Cluster-aware split: stratified group k-fold splitting grouped by the 30% sequence clusters.
5. Protein-family holdouts: predefined holdouts for major drug-target families, including protein kinases, G protein–coupled receptors, and nuclear hormone receptors. For these family holdouts, all training records belonging to any sequence cluster present in the held-out family test set were removed from the training partition to enforce cluster-disjointness.

A permutation test was additionally performed by shuffling labels within the training set while preserving feature distributions and grouping structure, followed by repeating the full tuning and evaluation workflow to establish a null baseline.

In addition to measuring true positive recall, recall@K_i_ was computed on a per-structure basis to assess within-structure ranking: for each protein chain *i* with *K*_*i*_ known positive pockets in the test set, pockets were ranked by predicted probability and the metric was defined as the fraction of those *K*_*i*_ positives recovered within the top *K*_*i*_ ranked pockets for that same chain, averaged across all test-set chains with *K*_*i*_ > 0.

#### Temporal and novelty-based external evaluation for GrASP comparison

To enable an unbiased comparison with GrASP^13^, whose model was published on July 26, 2023, we constructed a temporally and sequence-disjoint evaluation set. Specifically, we identified all PDB chains released after July 26, 2023, and mapped them to the global sequence-based clustering of PDB SEQRES entries (30% identity, MMseqs2).

From this pool of post-publication structures, we retained only those chains belonging to sequence clusters that were not observed in any PDB entry released prior to July 26, 2023. This ensured that the evaluation set comprised genuinely novel sequence families relative to the GrASP training period, preventing indirect sequence-level information leakage.

To avoid overrepresentation of homologous proteins, we selected one structure per novel sequence cluster. When multiple eligible chains existed within a cluster, a single representative was retained. This procedure resulted in a final evaluation set of 26 unique PDB chains spanning previously unseen sequence clusters. This temporally separated, cluster-disjoint dataset was used to benchmark PocketBagger against GrASP under a forward-looking, prospective-style evaluation scenario.

An instance of our PocketBagger model was then tuned and trained as outlined above using only structures available before July 26, 2023. The hold-out GrASP set was then predicted on. For this comparison, we evaluated GrASP on the 26-chain temporal/novelty holdout set and converted its atom-level predictions into a site-level score anchored to the experimentally observed ligand. For each held-out structure, we defined the GrASP “site” as the set of protein heavy atoms within 4 Å of the bound drug-like ligand, and summarized GrASP’s atom-level probabilities over that set using two aggregations: the mean probability and the maximum probability. Site-level recall was then computed using GrASP’s recommended classification threshold of 0.3, counting a site as recovered if either the mean-based score (mean ≥ 0.3) or max-based score (max ≥ 0.3) exceeded the threshold; we report recall under both definitions to capture conservative (mean) versus permissive (max) site calling.

#### Final production model and performance reporting

A final production PocketBagger model was trained using all available pocket records (positives and unlabeled) and the same Bayesian hyperparameter optimization strategy described above, including cluster-aware validation folds and average precision as the tuning metric. Because no independent label set exists for true negatives, we report the nested cross-validation recall and recall@K_i_ from the above evaluation regimes as the primary estimate of expected generalization performance for the production model. Additionally, a final isolation forest model was trained during nested cross-validation on the complete set of positives to be deployed alongside the production model.

#### Out-of-fold predictions and feature importance

To generate out-of-fold (OOF) predictions for downstream analyses, the production training set was split into five stratified group-aware folds using sequence clusters as groups. For each fold, a model was trained on the remaining folds with the final model’s hyperparameters and used to predict scores for the held-out fold; these predictions were concatenated to produce OOF scores for all pockets.

Feature importance was quantified using permutation importance computed on the held-out fold for each of the five folds, using average precision as the scoring function. Importances were aggregated across folds to obtain a stable estimate of global feature contributions under cluster-disjoint evaluation. To summarize feature redundancy and improve interpretability of importance estimates, we clustered pocket descriptors based on pairwise linear correlation structure. Specifically, we computed a pairwise linear feature–feature distance matrix as 1 – *R*^*2*^ (where *R* is the Pearson correlation between features across the training set) and applied agglomerative hierarchical clustering with average linkage on this precomputed distance matrix. The optimal threshold was found following a Bayesian optimization of the silhouette score, resulting in an optimal threshold of roughly 0.79 (corresponding to an *R*^*2*^ of 0.21) and a silhouette score of roughly 0.35 and a total of 11 clusters.

#### Positive Class Thresholding in Production

Because PocketBagger is trained in a positive–unlabeled setting, we supplemented model scores with several complementary, data-driven heuristics to determine where unlabeled pockets begin to look “positive” in production based on the above computed out of fold scores. First, we computed Tukey-fence thresholds on the distribution of predicted probabilities for the unlabeled pool, using upper fences at k=1.5 and k=3.0 to provide different stringency cutoffs for calling high-scoring unlabeled pockets positive. Second, we learned an isolation-forest score boundary to identify pockets that are statistically typical of the positive class. From the production isolation forest anomaly scores, a simple depth-1 classifier was fit to the resulting isolation scores to derive a single score threshold separating “positive-like” pockets from more anomalous ones. Third, we applied an Elkan & Noto (PU) class-prior estimation procedure by training a calibrated probabilistic classifier (isotonic calibration) to model *P*(*s* = 1 ∣ *x*), estimating *c* = 𝔼[*P*(*s* = 1 ∣ *x*) ∣ *y* = 1] on labeled positives, and converting this into an estimated positive class prior *π*over the full dataset. The resulting class prior was then used to derive a score cutoff corresponding to the top *π* fraction of pockets, providing a principled operating point for “likely positive” predictions. Taken together, these three approaches – unlabeled Tukey fences, positive-like isolation thresholds, and Elkan–Noto prior-based cutoffs – offer practical, auditable guidance for selecting prediction thresholds when deploying PocketBagger to score unlabeled pockets at scale.

## Acknowledgments

The authors would like to thank the canSAR technical team for their efforts in database management and data engineering. B.A-L. is a Cancer Prevention & Research Institute of Texas (CPRIT) Scholar in Cancer Research and is grateful for their support. She is funded under the Cancer Prevention and Research Institute of Texas (CPRIT) Established Investigator Award (RR210007). BA-L acknowledges additional funding from The Commonwealth Foundation, The Lyda Hill Foundation, the CRUK Drug Discovery Committee strategic award, NIHR (National Institute of Health Research) support to the Biomedical Research Centre at the Institute of Cancer Research, and the Royal Marsden NHS Foundation Trust.

## Funding

Cancer Prevention and Research Institute of Texas (CPRIT) Established Investigator Award (RR210007), (BA-L)

The Commonwealth Foundation, (BA-L)

The Lyda Hill Foundation, (BA-L)

## Author contributions

Conceptualization: PWG

Methodology: PWG

Investigation: AB, PDM, PWG

Funding acquisition: BA-L

Project administration: BA-L

Software: ILM, KMB, ZS, DSM

Supervision: BA-L, PWG

Writing – original draft: PWG, AB, KPR

Writing – review & editing: PWG, KPR, AB, BA-L

## Competing interests

B.A.-L. declares financial interest in Exscientia PLC, Recursion Pharmaceuticals Inc and AstraZeneca PLC. B.A.-L. is/has been a member of Scientific Advisory Boards and/or provided paid consultancy for the following: Astex Pharmaceuticals, AstraZeneca PLC, GSK PLC, Novo Nordisk, Sante Ventures. She is a lead on the MD Anderson Drug Discovery and Development Division which has commercial interest in target and drug discovery. She was a chair and member of the Scientific Advisory Board for Open Targets. She is chair of the Cancer Research UK Data Strategy Board and member of the CRUK Scientific Advisory Board. She is a member of the New York Genome Consortium Scientific Advisory Board. She is Director of non-profit Chemical Probes Portal. B.A.-L has been an employee of the ICR, which has a Rewards to Inventors scheme and a commercial interest in the development of cancer drug targets. P.G., A.B., K.R., and B.A.-L. are employees of UT MD Anderson Cancer Center which operates a reward to inventor scheme.

## Data and materials availability

The PU druggability classifier is deployed in canSAR which is freely available at https://cansar.ai. All source code is available at our GitHub repository: https://github.com/a3d3a.

## Supplementary materials

Included in separate Supplementary Materials.

## Notes

https://github.com/a3d3a

https://cansar.ai

