## Supplementary Text for "PocketBagger: Generalizable pocket druggability prediction via positive–unlabeled learning"

**Supplementary material for:** PocketBagger: Generalizable pocket druggability prediction via positive–unlabeled learning

**Authors:** Phillip W Gingrich^1^, Ansuman Biswas^1^, Ioan L Mica^2^, Kevin M Brammer^2^, Zhigang Shu^2^, David S Maxwell^1^, Kaitlyn P Russell^1^, and Bissan Al-lazikani^1,3*^

^1^Department of Genomic Medicine and the Institute for Data Science in Oncology, UT MD Anderson Cancer Center; Houston, Texas, USA

^2^Enterprise Development and Integration, UT MD Anderson Cancer Center; Houston, Texas, USA

^3^The Therapeutic Discovery Division; UT MD Anderson Cancer Center; Houston, Texas, USA

**Comparisons to other druggability predictors**

Direct comparisons of performance to other druggability predictors are challenging due to differences in the unit of prediction and evaluation protocols. Deep learning methods such as GrASP and P2Rank generate predictions at the atom or surface point level, which are subsequently clustered into predicted binding sites. Performance is then evaluated using distance-based criteria, measuring the distance between predicted site centers and either the nearest ligand heavy atom (DCA, as in GrASP) or the ligand center of mass (DCC, as in P2Rank), with distances < 4 Å considered successful. Both methods report recall-like metrics based on the top-N predicted sites, where N corresponds to the number of known ligands in each structure. This ranking-based evaluation makes Recall@Kᵢ a more directly comparable metric for PocketBagger.

Using this comparison, PocketBagger achieves a Recall@Kᵢ of 0.805 ± 0.059 under stringent cluster-aware splitting, which is comparable to the reported DCA/DCC top-N recall for GrASP (0.813) and P2Rank (0.812) on the HOLO4K dataset, and slightly exceeds the corresponding values reported for GrASP (0.775) and P2Rank (0.749) on the COACH420 dataset. We note that these comparisons are not strictly equivalent, as the data splits, ligand definitions, and evaluation protocols differ between methods. Nonetheless, these results suggest that a positive-unlabeled learning framework with an explicit emphasis on generalizability can achieve performance comparable to state-of-the-art deep learning approaches trained using standard supervised formulations.

**Pocket Descriptors**

Below are the definitions of features used as pocket descriptors.

ACC_BUR_VERT_RATIO: accessible/buried vertices ratio -> ENCLOSURE

ACC_RATIO: Accessible ratio

ACC_VERTICES: Accessible Vertices

ANDREWS_ENERGY: Inverse Andrews Energy

BETA_SHEET: % of residue in BETA sheet

BUR_VERTICES: buried Vertices

CONS_RATING: conservation rating -> average score on all spheres within 5A of average position

HP_RATIO: Hydrophobic/Polar Ratio

HB_ACCEPTOR: number of Hydrogen Bond Acceptor atoms

HB_BOTH: number of Hydrogen Bond acceptor and donor at the same time atoms

HB_DONOR: number of Hydrogen Bond donor atoms

HELIX: % of residue in APLHA helix

HOT_FRACTION: number of reactive groups in the pocket as a fraction of all group

LONG_AXIS: long axis value

LOOP: % of residue in a loop

MAX_DEPTH: Max depth value

MEAN_AXIS: Mean axis value

NRM_HYD_RATIO: Hydrophobic Ratio

NRM_POLAR_RATIO: Polar Ratio

PCA_X: Principle axis components value

PCA_Y: Principle axis components value

PCA_Z: Principle axis components value

POCKET_SIZE: Volume

TURN: % of residue in TURN

VOL_RATIO: Volume ratio

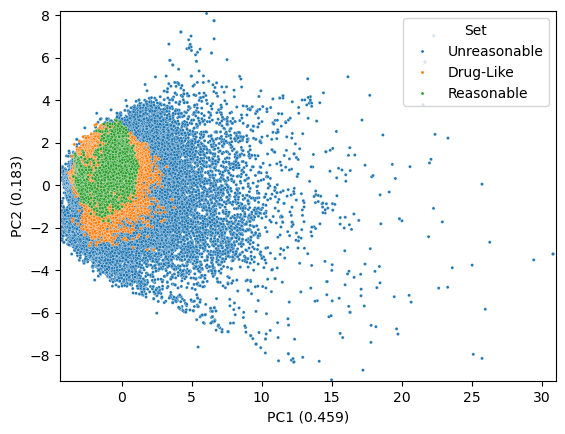

Figure S1 – PCA of “drug-like”, “reasonable”, and “unreasonable” compounds. Values in parentheses indicate the explained variance ratio for each component.

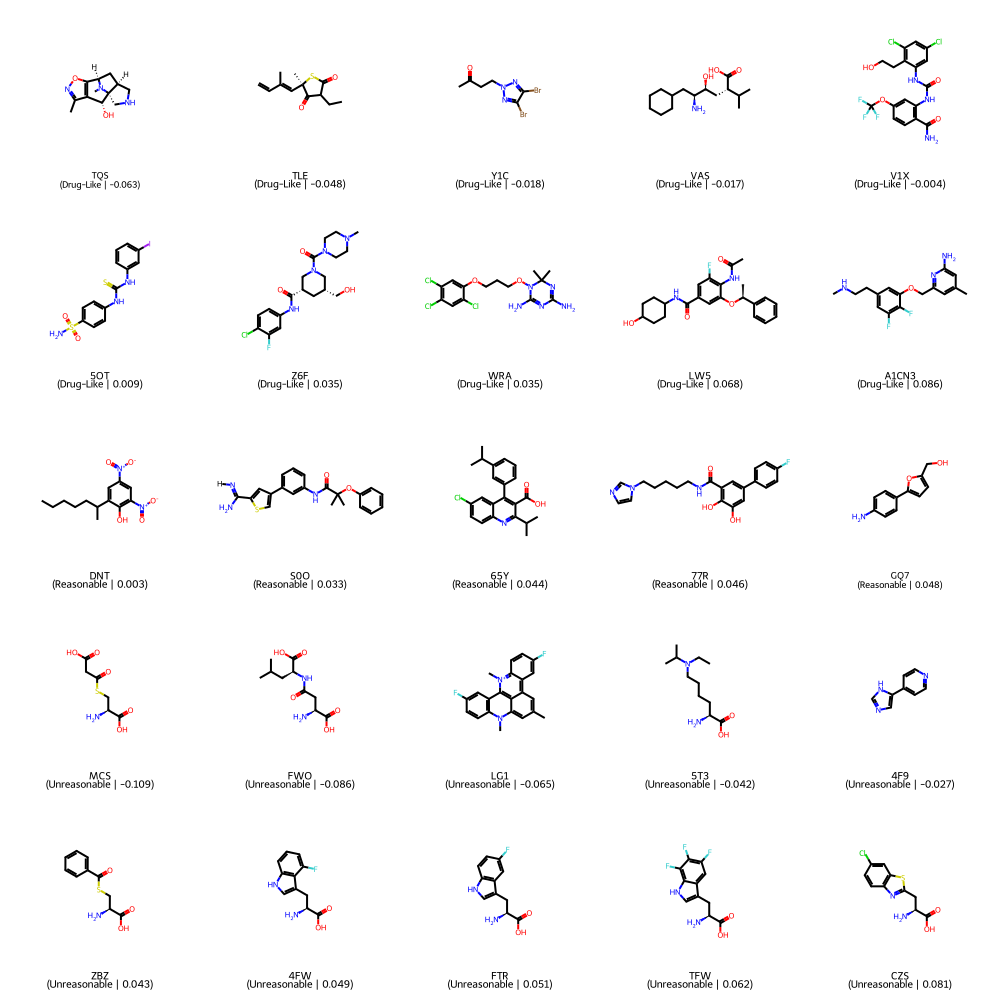

Figure S2 – Examples of “drug-like”, “reasonable’, and “unreasonable” HET records. Pockets occupied by “drug-like” and “reasonable” compounds were labeled as positive. The set and the record’s isolation forest anomaly score are presented in parentheses.

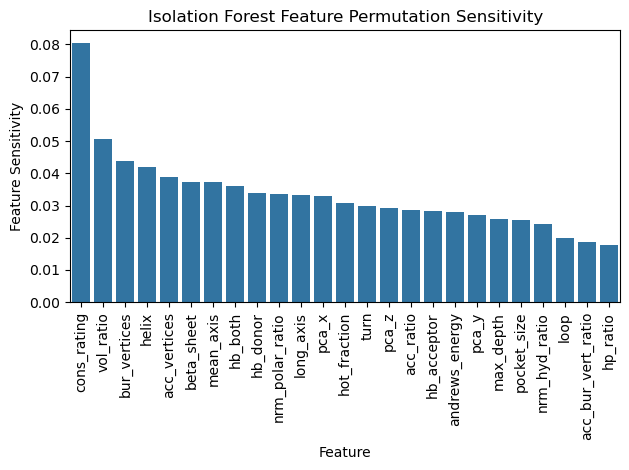

Figure S3 – Isolation Forest feature sensitivity results. High sensitivity indicates the associated feature was significant in scoring how anomalous pockets are for positives.

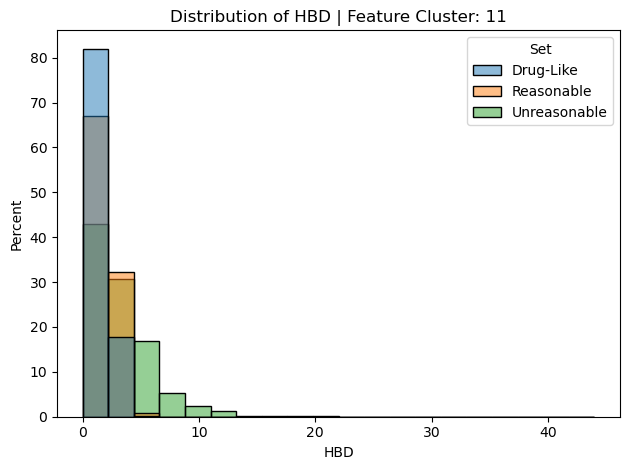

Figure S4 – Property distribution for HBD.

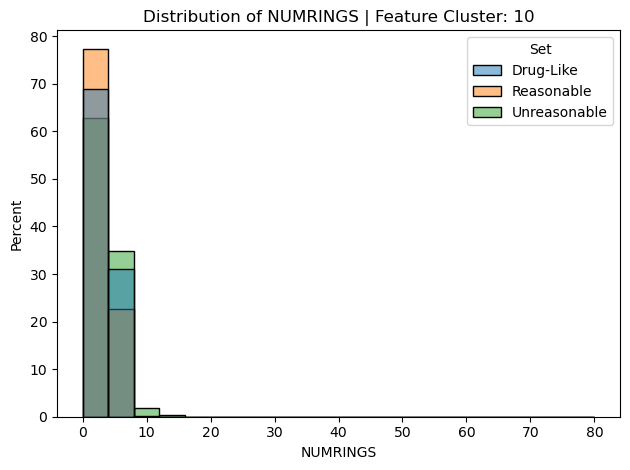

Figure S5 – Property distribution for NUMRINGS.

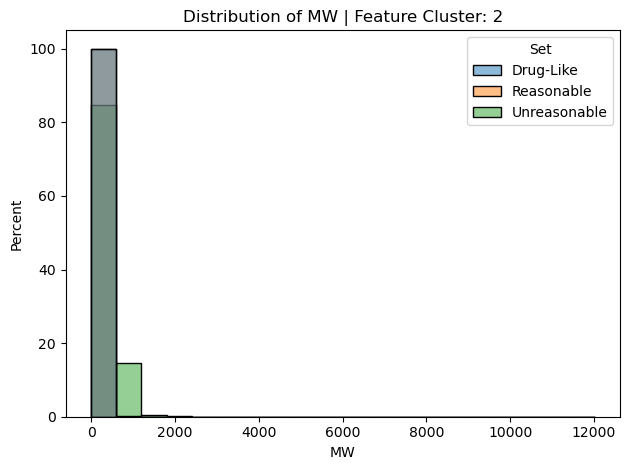

Figure S6 – Property distribution for MW.

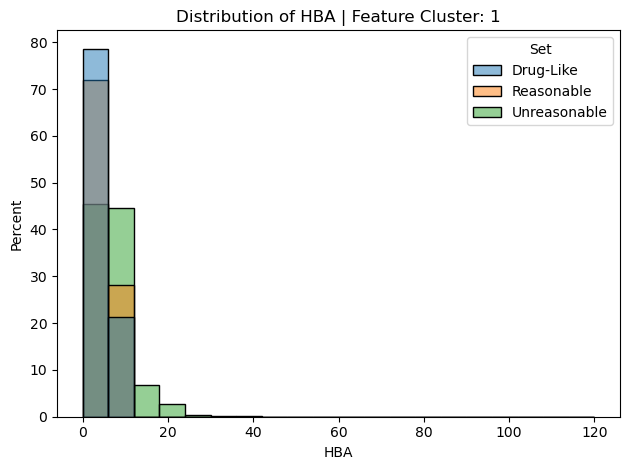

Figure S7 – Property distribution for HBA.

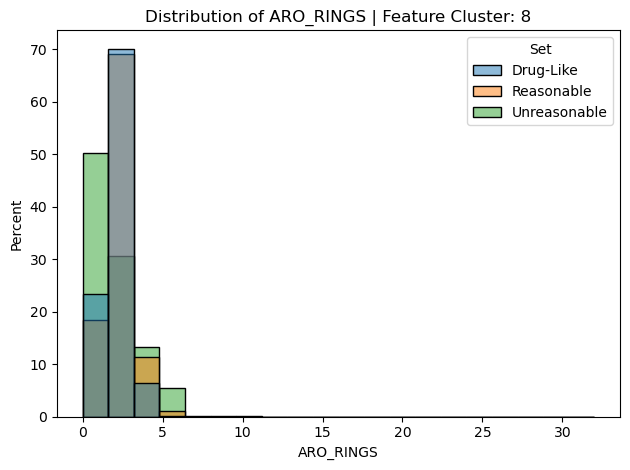

Figure S8 – Property distribution for ARO-RINGS.

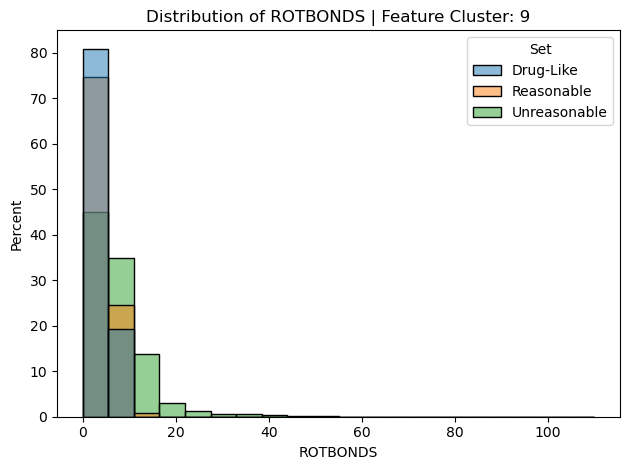

Figure S9 – Property distribution for ROTBONDS.

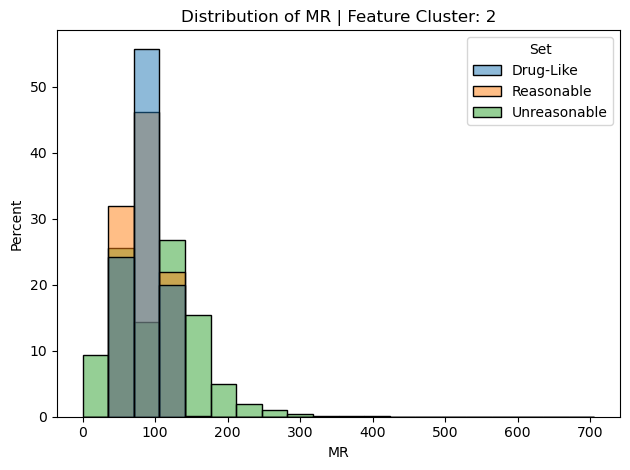

Figure S10 – Property distribution for MR.

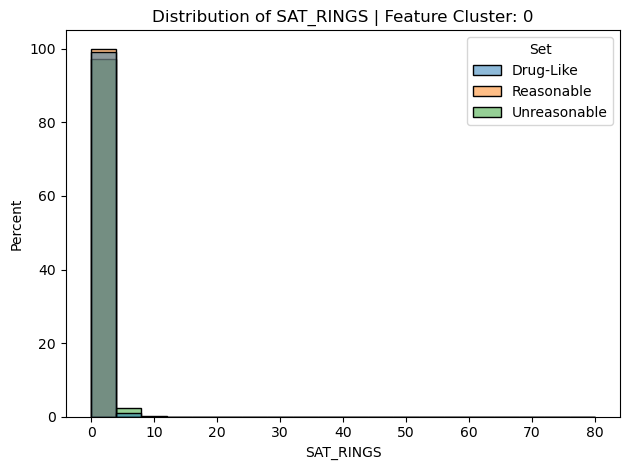

Figure S11 – Property distribution for SAT_RINGS.

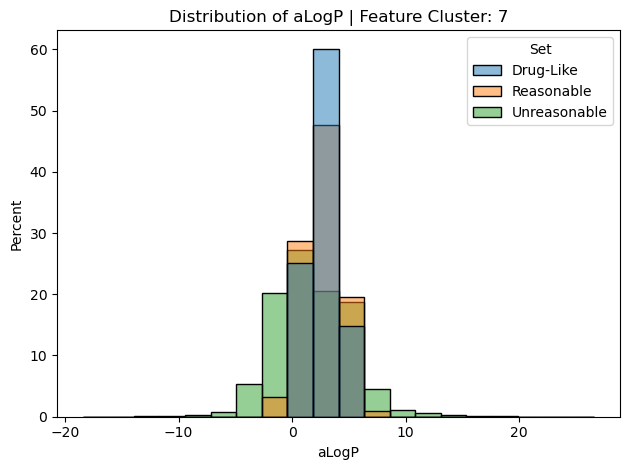

Figure S12 – Property distribution for aLogP.

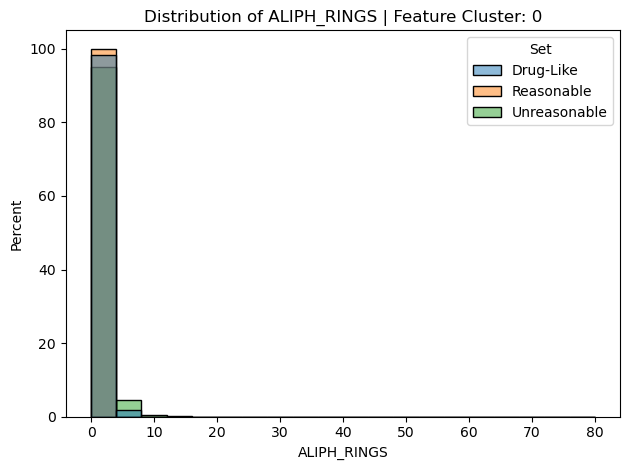

Figure S13 – Property distribution for ALIPH_RINGS.

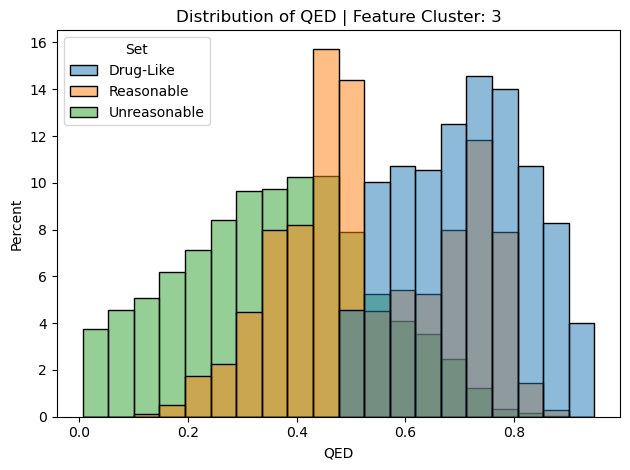

Figure S14 – Property distribution for QED.

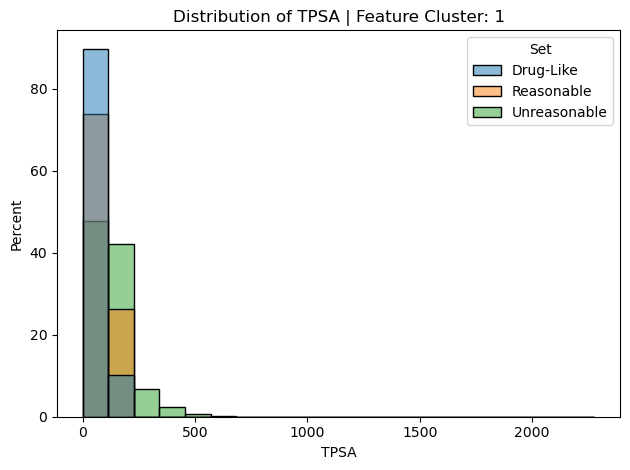

Figure S15 – Property distribution for TPSA.

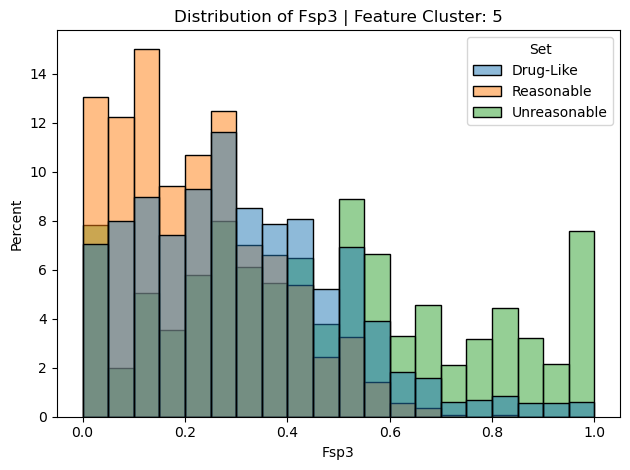

Figure S16 – Property distribution for Fsp3.

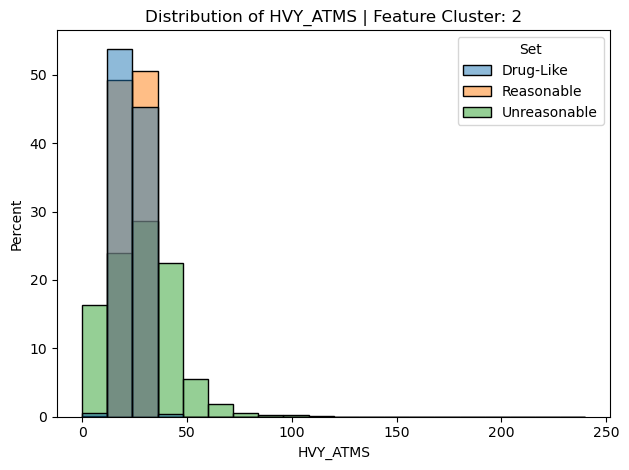

Figure S17 – Property distribution for HVY_ATMS.

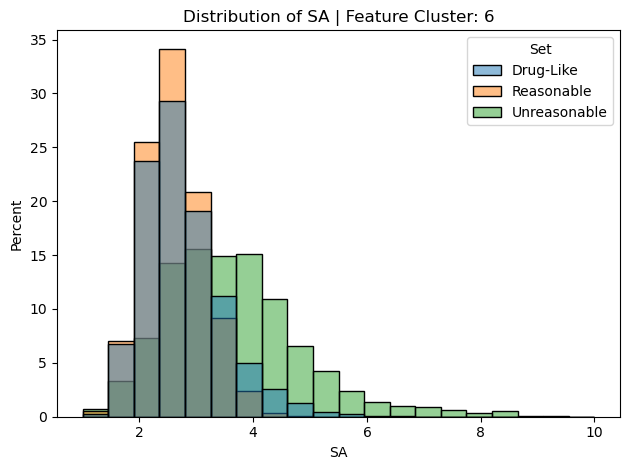

Figure S18 – Property distribution for SA.

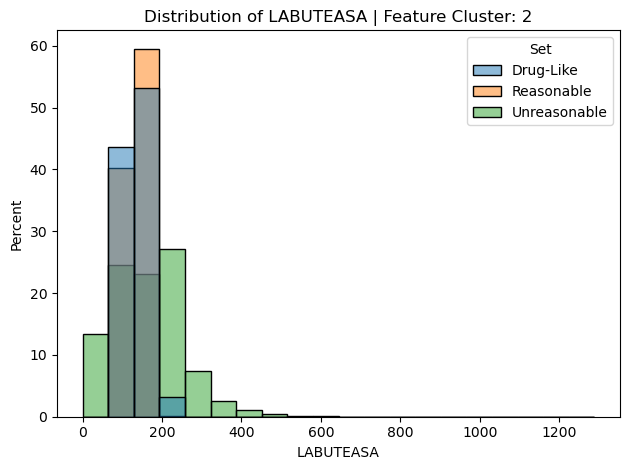

Figure S19 – Property distribution for LABUTEASA.

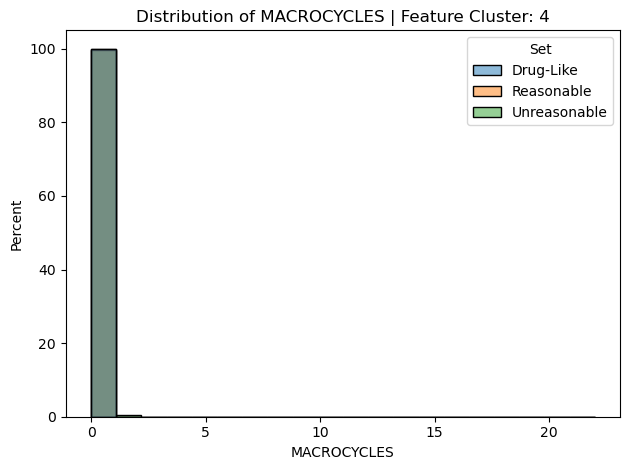

Figure S20 – Property distribution for MACROCYCLES.
